# Wavelet analysis of dual-fMRI-hyperscanning reveals cooperation and communication dependent effects on inter-brain neuronal coherence

**DOI:** 10.1101/2025.03.27.645650

**Authors:** Rik Sijben, Robert Friedmann, Lucia Hernandez-Pena, Rea Rodriguez-Raecke

**Affiliations:** Brain Imaging Facility, Interdisciplinary Center for Clinical Research (IZKF), RWTH Aachen University, Aachen, Germany; Department of Psychiatry, Psychotherapy and Psychosomatics, Faculty of Medicine, RWTH Aachen University, Aachen, Germany; JARA – Translational Brain Medicine, Aachen, Germany

**Keywords:** functional magnetic resonance imaging, hyperscanning, wavelet transform coherence, social interaction, cooperation

## Abstract

Hyperscanning has allowed neuroscience to expand investigations into neuronal activation during social interactions. Rather than analyzing how a single brain responds, we can compare interactions and even synchronization between multiple actors in varying situations. This technique is commonly employed using functional near infrared spectroscopy (fNIRS). Specifically, social cooperation and competition have been thoroughly investigated using this approach. While functional magnetic resonance imaging (fMRI)-based hyperscanning is becoming more prevalent, a link to this fNIRS-based foundation is missing. We here use a dual-fMRI-hyperscanning setup and an established task to investigate neuronal coherence during social cooperative and competitive tasks. Wavelet transform coherence (WTC) allows us to explore task-specific frequency bands of interest of non-stationary neuronal activation signals of paired participants (n=60). We show that cooperation, compared to a control task, increases inter-brain neuronal coherence in regions associated with social interaction and the theory of mind (ToM) network. Verbal communication prior to the task expands this coherence to different regions of this network, including middle and superior temporal gyrus. This spatial shift suggests additional implementations of the ToM network depending on the cooperation approach taken by the participants. Our findings both support and expand on results by previous fNIRS-based studies and show that WTC is an effective way to model fMRI-based neuronal synchronization; thereby closing the gap between two popular hyperscanning methodologies.

**Significance Statement:** Within social neuroscience a strong basis exists for fNIRS-based hyperscanning. In the past years fMRI-based hyperscanning has increased in popularity, yet the basis for these projects seems to develop independent of the existing background. The current study aims to connect these previous findings to the possibilities offered by fMRI, allowing both fields to benefit from the other’s advantages when determining optimal research paradigms or analyses.

## Introduction

Social cooperation relies on the interaction between individuals who acknowledge each other’s identity. A topic of interest for social neuroscience is to determine whether or how the neural correlates of this behavior might interact. Investigating this can, however, be challenging as neuroscientific methodologies are generally optimized to investigate a single participant. Yet neuronal activation of a single volunteer cannot sufficiently explain a process which relies on the interaction of multiple individuals. Hyperscanning (HS), the act of simultaneously measuring neural activation of two or more participants, is a relatively novel approach, slowly gaining traction since the term was coined by Montague et al. ^1^. The technique’s popularity seems mostly driven by electroencephalographic (EEG) and functional near-infrared spectroscopy (fNIRS) research. Though Montague initiated the technique using a functional magnetic resonance imaging (fMRI) setup, HS-fMRI studies currently make up less than 10% of the field’s literature. This sparse representation can be attributed to the technical and financial burden of operating and synchronizing two fMRI scanners, whereas EEG and fNIRS HS paradigms do not even necessitate a second hardware setup ^2^. An important contribution to the field was made by Cui et al.^2^ who utilized an HS-fNIRS approach to investigate synchronization of oxygenated hemoglobin concentrations (HbO) in the frontal cortices of participant dyads that performed a cooperation task. In this task, participants were instructed to respond to a color-changing visual cue by simply pressing a button. The task objective could easily be switched from “cooperation”, during which participants completed the task by responding simultaneously within a margin of error, to “competitive”, during which participants were instructed to respond as fast as possible. A control task “single” was performed by letting each participant perform the task alone. Two measurement channels, corresponding to right superior frontal cortex, showed significantly higher coherence in HbO levels during cooperation tasks compared to the other conditions.

Cui et al. ^2^ introduced a straightforward, easily customizable task and showed the feasibility of wavelet transform coherence (WTC) as a measure of inter-brain neuronal synchronization. WTC is a time-frequency decomposition method, that was initially used for meteorological ^3,4^ and geophysical analyses^5^, and later applied to neuronal signals ^6^. Briefly, (continuous) wavelet transform convolves a time series with a pre-defined wavelet which, in the case of the analytical Morlet wavelet, includes a specific angular frequency and Gaussian envelope. This decomposes the underlying signal in the time-frequency domain with an imaginary part indicating phase. Coherence then describes the phase-locked similarities within this domain between two signals of uniform length. Figure 1 visualizes an example wavelet, continuous wavelet transform, and WTC of two non-stationary signals. For a more in-depth description of WTC see Torrence and Compo ^3^ or Chang and Glover^6^ for its applications in neuronal signal analysis.

**Figure 1.**
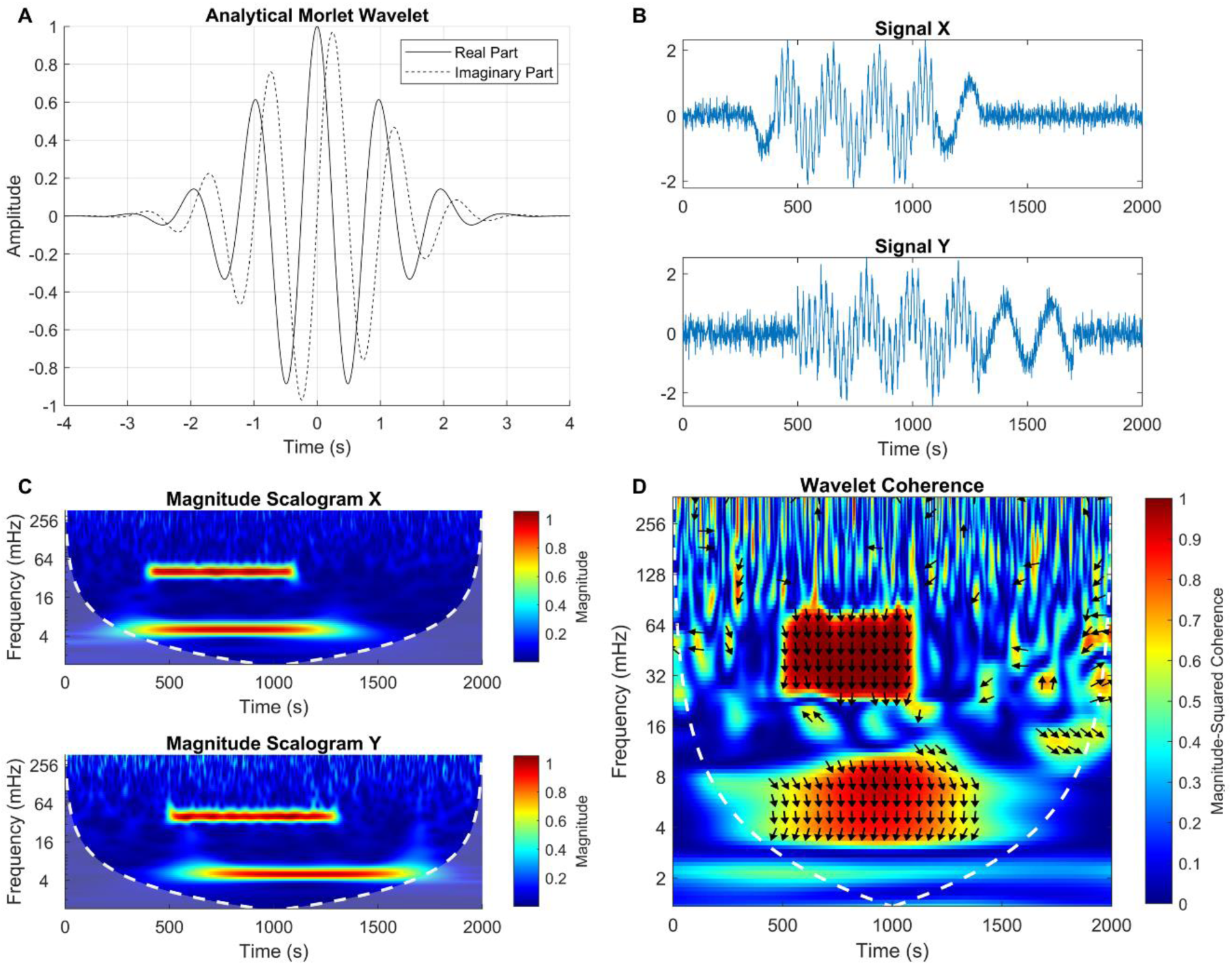
Brief overview of wavelet transform coherence (WTC). **A)** Example analytical Morlet wavelet (ω = 2π, σ = 1). **B)** Example signals consisting of partially overlapping sine waves (X) or cosine waves (Y) of 5 and 40 mHz, contaminated with Gaussian noise. **C)** Scalogram visualizing the magnitude of specific frequencies in signals X and Y across time, generated by a continuous wavelet transform of the signals with an automatically scaled analytical Morlet wavelet. The color bar corresponds to relative power. Dotted white lines indicate the cone of influence boundary, beyond which edge effects are too influential to meaningfully interpret the data. **D)** Wavelet coherence of signals X and Y. The heatmap visualizes the magnitude squared coherence at different frequences across time. Regions of warm colors indicate strongly phase-locked coherence between the two signals. Two clear synchronized clusters are seen corresponding to the low and high frequency components of the signals. The phase shift between the synchronized components is indicated by the direction of the arrows; a vertical downwards pointing arrow indicates a phase shift of π/2, prominent in the example as X and Y contain strong sine and cosine components, respectively.

WTC offers several advantages for the analysis of HS-induced neuronal activation patterns. Primarily, its consideration of non-stationary signals fits the experimental design of psychological studies as we expect synchronization to vary as the task changes. Second, the separation in frequency scales allows one to limit the analysis to task-relevant activation patterns, filtering out frequencies of no-interest or simply increasing statistical power. Finally, by allowing phase-shifts between the two signals, models of neuronal activation can reflect reality more closely.

The paradigm designed by Cui et al. ^2^ has been implemented in several similar fNIRS projects. Cheng et al. ^7^ and Baker et al. ^8^, for example, both showed gender pairing effects within the dyads. Though Cheng et al. ^7^ reported no significant task related coherence differences after appropriate corrections were applied, coherence differences were found for male/female dyads for several channels corresponding to orbitofrontal cortex and dorsolateral prefrontal cortex. Baker et al. ^8^ added an additional target by adding a second optode patch covering the right temporal lobe in addition to a right inferior prefrontal one. Using this setup, they reported that male/male dyads showed significantly higher coherence levels in right prefrontal cortex during cooperation compared to male/female or female/female dyads. Right temporal regions, however, showed higher coherence during cooperation for female/female dyads. Pan et al. ^9^ further expanded these findings by differentiating familiarity levels of the dyads. By categorizing participants as “strangers”, “friends”, and “lovers”, they were able to show that increased familiarity not only improved performance on the cooperation task, but also increased synchronization of right superior frontal cortex. Child-friendly variants of this paradigm were also implemented to investigate autism spectrum disorders ^10^ and mother-child dyads ^11^. When participants attempt to synchronize their responses, they are focusing on a mutual goal. Therefore, previous studies^2,7–9^ have described this as social cooperation. As participants are often not allowed, or even able, to communicate, one could argue that attempting to achieve this mutual objective could just as well be described as coordination. As such, caution should be taken when comparing the task to other forms of cooperation which involve direct social interaction. For consistency with previous work, we will continue using the term “cooperation” in the current paper.

Compared to fMRI, fNIRS offers several straightforward advantages including ease of use and relatively high sampling rate. Above all, fNIRS studies can obtain a high level of ecological validity as mobile setups such as the NIRSport2 (NIRC Medical Technologies, nirx.net/nirsport) enable realistic research environments, especially in comparison to fMRI. The disadvantages of fNIRS, however, become apparent when one attempts to localize the observed coherence. HS-fNIRS projects often limit themselves to a single region of interest, usually a broad sub-region of the frontal cortex, or generalize over another large region ^8^. Employing an fMRI setup allows for a whole-brain analysis, potentially revealing neuronal synchronization in regions that have not yet been analyzed or cannot be investigated using fNIRS due to their anatomical location. Additionally, the spatial specificity offered by fMRI allows us to specify the responsible regions in more detail. Inter-brain coupling can be, and has been, analyzed and interpreted using numerous methodological approaches. A recent review ^12^ exploring hyperscanning studies published between 2000 and 2022 describes the 27 different methodologies used across 4 modalities in 215 studies. In the current work we aim to close the gap between two of these modalities by employing the most common methodology (WTC). Our choice of this approach should not be seen as a value statement regarding the other methodologies.

The development of fNIRS and fMRI HS platforms is driven primarily by social neuroscience, with cooperative behavior being a particular point of interest. Remarkably, most studies using variants of the task by Cui et al. ^2^ choose to limit this social cooperation by instructing participants to refrain from verbal communication or even adding a barrier between participants to avoid any form of contact ^9^. While this choice improves consistency of the paradigm, it limits interpretation and generalizability of the findings. As the synchronization found by previous studies is, in part, attributed to the (indirect) social interaction required for the task, the question arises of how verbal communication would influence this effect. We suggest two possible mechanisms driving neuronal synchronization which could be unraveled by introducing verbal communication: Our first hypothesis argues that the behavior of two participants trying to match each other’s actions is indeed a form of social cooperation. If this is the case, then we would expect that the introduction of verbal communication further enhances this social component, thereby increasing neuronal synchronization in the same regions seen during the cooperative task without communication. Our alternative hypothesis argues that it is in fact the act of separating the participants which drives neuronal synchronization. Successful completion of the task required the participants to anticipate each other’s actions (responding fast or responding slow). As no contact is possible between participants they must rely on their theory of mind (ToM)^13–16^, driven by the participant’s previous actions. If this hypothesis is correct, then allowing the participants to directly communicate their intentions lowers this need to anticipate behavior and neuronal synchronization is likely to be either reduced or shifted to other regions relevant for communication.

For the current study we set up a dual-fMRI-HS platform (Figure 2A) to investigate neuronal coherence within male dyads while they complete a two-person interactive computer task based on the design by Cui et al. ^2^. An additional experimental task is added to this paradigm; during this task, verbal communication is enabled through the use of active noise cancellation headphone and microphone systems at both MRI scanners. With this setup we expect to reproduce previous fNIRS based findings, showing cooperation dependent effects on neuronal coherence. We then expect to expand these findings to broader neural networks related to social interaction. Finally, we expect that the addition of verbal communication will change coherence patterns compared to mute cooperation. The form in which these changes take place can inform us about the underlying process driving the coherence. By employing a similar task and analysis, the current study aims to serve as a link between the existing HS-fNIRS research and the currently expanding field of HS-fMRI.

**Figure 2.**
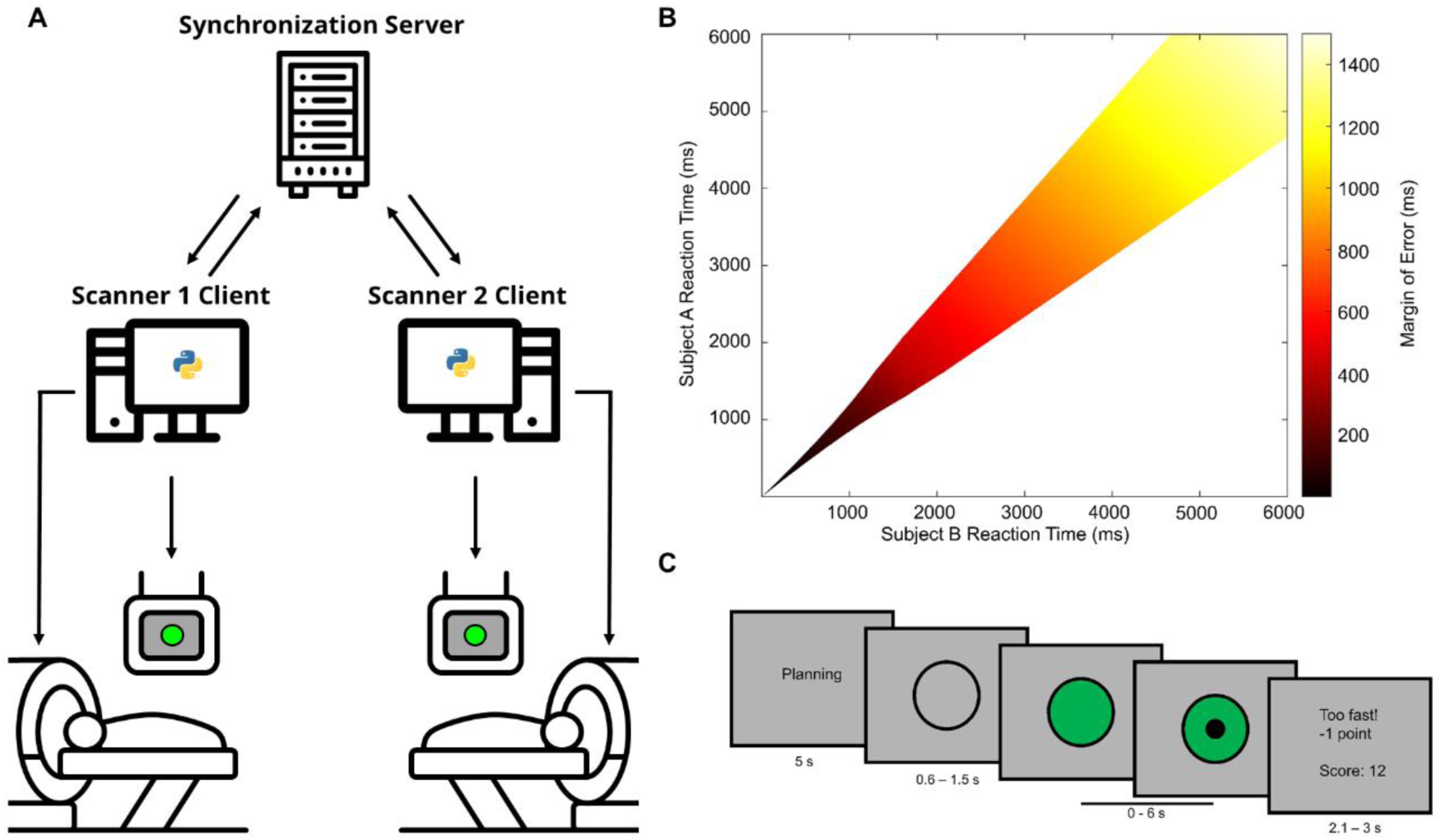
**A)** Dual-fMRI hyperscanning setup. Scanner clients send paradigm relevant information and are synchronized over a locally hosted server. fMRI measurements were started simultaneously through a trigger from the respective scanner clients. **B)** Heatmap depicting the variable margin of error used to determine simultaneous responses during the cooperation and communication tasks. Response times falling within the colored space were considered simultaneous. Fast responses were discouraged by decreasing the margin of error exponentially for values below 1600 ms. **C)** Example trial of the behavioral task. During the first 5 seconds participants were encouraged to think about how to respond to the coming stimulus. The gray circle had a variable duration after which it is colored green. Participants pressed a button following task specific instructions. A black dot was visible while waiting for the other participant. Feedback was presented after each trial.

## Materials and Methods

### Participants

A total of 66 male volunteers aged 18 to 40 (Median = 22 years) participated in the study. Participants were recruited through flyers spread around university buildings, student housing, and train stations around the city of Aachen, Germany. Volunteers were included in the study only if they were aged between 18 and 40, were physically and mentally healthy, and were free of MRI contra-indications (e.g. metal implants, claustrophobia). Forty-eight volunteers signed up in pairs, the remaining 18 were randomly paired with another volunteer who spoke the same language. All included volunteers were able to read and understand the German or English instructions. During the communication task, any preferred language was allowed.

Previous studies showed that neuronal coherence in the context of cooperation presents a sex and/or gender effect ^7,8^. Considering the high operating costs of fMRI-HS, we were forced to constrain the total required sample size. As such, only male volunteers were recruited as previous studies reported larger effect sizes for male participants^7,8^. Volunteers received a digital copy of the informed consent forms several days prior to their participation to ensure ample time to read and ask questions. On the day of the study, the volunteers signed the forms, providing their written informed consent. After completion of the experiment, participants were asked if, at any point during the study, they doubted whether the interaction with the other participant was real. After answering they were ensured that no manipulation took place and were given the chance to ask detailed questions about the research question and scientific purpose of the tasks they performed. The study conformed to the ethical principles of the World Medical Association’s declaration of Helsinki^17^. The study was approved by the ethical committee of the medical faculty of the RWTH Aachen University (EK 387/20) and Center for Translational & Clinical Research (CTC-A-# 20-407) and was registered in the German Clinical Trials Register (DRKS00030177) of the German Federal Institute for Drugs and Medical Devices (BfArM).

### Behavioral task & Procedure

Participants simultaneously performed a button pressing computer task (Figure 2C) similar to that employed by previous fNIRS studies ^2,7–11^. The task was programmed and run using PsychoPy3 ^18^ and was available in English and German. Participants indicated their preference for the language prior to participating. Each trial of this task was initiated by a 5-second “planning” phase during which the word “*Planning”/”Planung”* was shown on screen. Participants were instructed to use this time to consider how fast they would like to respond to the stimulus in the upcoming trial. After this, an empty black circle was presented. After a variable pseudorandomized delay, averaging 1050 ms, the center of this circle changed to a green color. The participants were instructed to press a button after this color change took place. After both participants pressed the button, or after a time-out (6 seconds), the participants received written feedback regarding their performance. The duration of this feedback varied, averaging 2550 ms, and was synced with the variable delay to ensure consistent trial durations. Participants were awarded a point when the specific task was completed successfully (i.e., simultaneous response, responding faster than opponent, responding at all in solo tasks) and were deducted a point when the task was failed (i.e., exceeding cooperation margin of error or responding slower than opponent). Additional points were deducted when the task was performed incorrectly (i.e., time-out or responding before the color change).

Trials were presented in different blocks; each block was preceded by an instruction screen reminding the participant about the specific goal for that task. These instructions were given prior to entering the MRI scanner as well. For the “Cooperation” block, participants were instructed to press the button simultaneously. For this task, the margin of error was determined by a two-stage computation: If the participant’s response time was below 1.6 seconds, it was adjusted through: 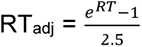. Following this, the margin of error for determination of synchronous responses was computed as 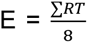 (Figure 2B). The adjustment for fast response times was implemented to discourage responding too fast, thereby increasing task difficulty.

The “Communication” block was predominantly similar to the cooperation block. They differed only in that participants were able to verbally communicate with one another during the planning phase. The on-screen instruction during this phase stated *“Planning. Communication possible.“/”Planung. Kommunikation möglich.”*. Note that the participants were no longer able to communicate once the circle was visible. Communication was enabled and disabled by automatically muting and unmuting the voice-chat software (Mumble ^19^) at the start and end of each planning phase using a custom python script. The objective of this task was still to press the button simultaneously, and the margin of error computation was identical to the cooperation task.

For the “Competition” block, participants received a different instruction; the objective was now to respond faster than the other participant, but still only after the color changed. No margin of error was applied here, the fastest responding participant was awarded a point each trial while the slower participant was deducted a point. This task generated a social cognitive component in that participants directly competed against each other, and the outcome was dependent on the other’s performance. However, the objective of the task itself was not affected by the behavior of the other participant; the goal was always based entirely on the stimulus, independent of the performance of the other participant.

The “Solo” and “Watch” tasks effectively functioned as paired control conditions. For the solo task, participant was instructed that his response time did not matter; as long as the participant pressed the button after the color change and within the 6-second time-out limit, a point was awarded. While one participant completed the solo task, the other half of the dyad performed the watch task. During this task, the participant observed the same visual stimuli as the other participant, and he was instructed to spectate the otheŕs performance. The spectating participant could not press a button and was not awarded points based on the other’s performance. This task was developed to remove the cognitive social component as neither the task, the result, nor the motivation for the task was affected by the other participant.

Participants completed each of the five blocks in sequence with short breaks in between. After the first run, containing each of the 5 tasks, participants were given a 5-minute break without a behavioral task. During this time a structural image of their brain was obtained. After this, the participants performed a second run during which each condition was presented a second time. The order of the solo/watch tasks was reversed for the second run. Task order was not randomized between dyads.

### Hyperscanning and synchronization

Precise synchronization of both computer tasks and MRI scanners is imperative for HS research. A server was hosted on a virtual machine at the Brain Imaging Facility of the University Hospital RWTH Aachen. This server was established using the socket module in a custom script using Python2.7. The task controlling computers (clients), located in the control rooms of both MRI scanners, communicated with this server through a TCP/IP network connection over the institute’s secure high-speed gigabit network. To improve accuracy, response times were calculated locally per client. These response times were then sent to the server in addition to a block identifier. Once the server received response times for each client the values were processed in a manner dependent on the accompanying block identifier and the appropriate feedback was replied to the clients. Based on this message the clients updated the respective scores and displayed the matching feedback. During the solo/watch blocks, the client engaged in the watch block generated an automatic response signal at the start of the trial as this participant could not press any buttons. The effect of client-server latency was dismissed as irrelevant as communication occurred at a latency of less than 4 milliseconds, well below the refresh rate of the displays used to run the task.

The clients running the computer tasks were simultaneously started by a signal from the server. Additionally, the task was set up to synchronize at every trial. This ensured that any sudden peaks in latency or even temporary loss of connection would not cause the two tasks to de-sync. No connectivity issues occurred during any of the measurements. To synchronize functional measurements, MRI sequences were set up to wait for user input before starting. As part of the synchronized (as described above) experimental task, PsychoPy generated output signals over a parallel port. These signals were then transformed by Arduino Uno^20^ micro-controllers to emulate user input (“Enter” key-press) to the scanner console, synchronizing the MRI scanners.

### Devices

Participantś communication during the MRI measurements was enabled by the OptoActive-II ANC headset and FOMRI-III optical noise cancelling microphone (Optoacoustics Ltd., Israel), implemented at both scanners. Proper noise cancelling is of vital importance for dyad communication during fMRI HS as each participant will hear not only the noise of their own scanner, but also any scanner noise picked up by the microphone of the paired participant. In addition to wearing the ANC headset, participants wore earplugs as the noise cancelling was only active during functional MRI sequences.

Participant communication was established using the open-source voice chat program Mumble ^19^. This specific program was chosen because it allowed us to self-host a server within the department, avoiding third-party entities, ensuring that data privacy was adhered to. Mumble ran on the same computer as the behavioral task. Computer audio was played directly to the participant’s headset and the microphone signal was selected as audio input within Mumble. By default, both participants were “deafened”, muting both microphone and audio. Both participants were automatically unmuted at the start of each planning phase during the communication task. At the end of this phase, they were automatically muted again. In addition to the visual instruction, an audible “ding” informed the participants when they were muted or unmuted. Communication between the participants was logged using the software’s built-in recording function. Audio quality and ANC functioning were verified at the start of each session using a test measurement. Audio levels were adjusted per dyad to ensure clear communication.

Stimuli were presented on MRI-compatible displays positioned behind the scanner bore. The displays were viewed through a mirror positioned above the participant’s eyes at an angle of 45°. scanner rooms were equipped with a 32 inch “BOLDscreen” (Cambridge Research Systems Ltd, Rochester, England). For one scanner the display offered a resolution of 1920 by 1200 pixels, while the other display offered a resolution of 1920 by 1080 pixels. Presentation size of visual stimuli was absolute and unaffected by pixel count of the displays. Participants performed the task using custom-built optical keypads positioned under their right hand.

### Magnetic Resonance Imaging

The study was conducted using two equivalent Siemens 3 Tesla whole body MRI scanners running syngo MR D13D; one Magnetom Prisma and one Magnetom Prisma Fit. B0 field strength and accordingly ^1^H-imaging frequency of the two scanners were compared and found to be within .05% of each other. All images were obtained using 20-channel head coils.

High-resolution structural images were obtained using a T1 weighted Magnetization Prepared Rapid Gradient Echo (MPRAGE) sequence, using the following parameters: repetition time (TR) = 2000 ms, echo time (TE) = 3.03 ms, slice gap = 0.5 mm, flip angle (FA) = 9°, matrix size = 256 x 256. The sequence yields 176 slices with a resolution of 1.0 mm isotropic in an acquisition time (TA) of 4 min. 16 s.

To optimize the analysis of signal coherence, temporal resolution of the functional data was of utmost importance. Therefore, functional images were obtained using a T2* weighted multi-band accelerated echo planar imaging (EPI) sequence using the following parameters: TR = 1500 ms, TE = 34 ms, no slice gap, FA = 70°, matrix size = 86x86, voxel size = 2.23 mm isotropic. The sequence yielded 57 slices per volume obtained in an interleaved order with a multi-band acceleration factor of 3. For the data sets included in the analysis, a mean of 775 (standard deviation = 23.1) volumes were obtained per run in a mean TA of 19 min. 23 s.

The B0 field was mapped using a double-echo gradient echo field map using the following parameters: TR = 526 ms, TEshort = 4.8 ms, TElong = 7.33 ms, no slice gap, FA = 60°, matrix size = 64x64, yielding 50 slices with a resolution of 3.0 mm isotropic.

## Analysis

### Pre-processing

MRI data of 3 dyads was excluded from the analysis due to interrupted measurements or structural abnormalities. Images were pre-processed using MATLAB 2023a ^21^, running SPM12 (v7771) ^22^. Prior to any analysis, the first 4 functional volumes acquired each run were discarded to ensure a steady-state magnetization was reached. Then the B0 field map, visualizing B0 inhomogeneities, was used to generate a voxel displacement map employed to correct distortion artefacts. Subject motion was corrected independently per run by realigning the images in a two-step procedure. Functional images were registered and resliced to each subject’s structural volume, aligning the images while benefiting from the improved level of detail offered by the high resolution MPRAGE. The structural images were then normalized to MNI-152 space and the corresponding transformation matrices were applied to the now aligned functional images, reslicing to a resolution of 2 mm isotropic. Finally, functional images were smoothed with a kernel size of 8 x 8 x 8 mm at full width at half maximum.

### General Linear Model

Pre-processed functional images of both runs were fed to a single general linear model (GLM) per subject; the different runs were modeled as such. Regressors of interest were generated by convolving stimulus specific boxcar functions with the canonical hemodynamic response function as implemented in SPM12. Task specific boxcar functions, corresponding to the 5 behavioral conditions, differentiating based on accuracy when possible, were generated based on the start of each trial. For this purpose, the trials ended when the subject pressed a button. Visual feedback presented to the subjects was captured in a single overarching regressor per run as condition specific variations in feedback processing were not of interest to us. Similarly, two regressors were added to capture the written instructions presented to the subject at the start of each block and the preceding rest blocks. Finally, per run 7 nuisance regressors were added, 6 capturing corrected subjected motion and 1 constant value. A 3.9 mHz (256 s period) high-pass filter was applied to remove scanner drift. An effects-of-interest F-contrast was generated for mask-based data extraction.

### Signal extraction and wavelet transform coherence computation

Analysis of wavelet transform coherence (WTC) in the context of fMRI has primarily focused on the variance of intra- or cross-network coherence ^6,23–27^. Network wide activation patterns and their coherence were distinguished as informative tools for classification of various complex heterogenous disorders ^23,25,27^. The aim of the current study, however, was to specify singular regions exhibiting inter-brain coherence. A number of atlases were considered for region-of-interest (ROI) selection ^28–31^. The Schaefer parcellations ^31^ are based on resting state fMRI data of 1489 healthy young adults from the Brain Genomics Superstruct Project ^32^ data set. Atlases with lower parcellation counts tend to generalize frontal cortices as several large regions and are unable to capture their functional heterogeneity. Therefore, we opted for the Schaefer-400 parcellation as it offers a good trade-off, creating usable ROIs while maintaining discernibility of different functional regions in frontal cortices. Anatomical descriptions of the different ROIs are assigned using the Harvard-Oxford cortical structural atlas ^29^.

First eigenvariates of the ROI’s blood oxygenation level dependent (BOLD) time courses were extracted per session using the *voi* command as implemented in SPM12. Signals were adjusted by effects-of-interest to eliminate nuisance factor related components from the signal. Further denoising was not performed. Data was extracted from 400 binary ROIs generated from the Schaefer-400 parcellation ^31^. To avoid sampling voxels with missing data, e.g. due to out-of-brain voxels or distortion artifacts, ROIs were masked by subject specific F-maps of the effects-of-interest contrast, thresholded at p < 0.9. ROIs were not selected nor masked based on statistically significant task effects to avoid circular analyses. Due to exact FOV positioning of the imaging sequence and distortion artifacts, data extraction was not possible for some ROIs for some dyads. ROIs suffering more than 33% missing data were excluded from the analysis entirely. 5 ROIs fulfilled this criterion (#28, #70, #207, #265, and #276 of the Shaefer-400 parcellation, matched to Kong-17 network order ^33^), corresponding to different parts of the rectal gyrus, medial orbitofrontal gyrus, and left anterior inferior temporal gyrus. Each of these ROIs is located in a region sensitive to susceptibility artifacts.

Once the signals Sa and Sb of a dyad were extracted, the magnitude-squared wavelet coherence was computed using the *wcoherence* function as implemented in MATLAB 2023a. Wavelet transforms wta and wtb of the signals were computed using de default analytic Morlet wavelet. Though an argument can be made for the generalized Morse wavelet ^11,34^, we prioritized the similarity to equivalent fNIRS studies ^2,7–9^. The frequency band for the transformation was set from 3.9 mHz to 289.4 mHz which yielded 75 scales generated at 12 voices per octave. The lower frequency limit matched the high-pass filter, and the upper frequency limit was based on the durations of the shortest measurement to ensure that a consistent number of scales (n=75) was generated despite the total number of functional volumes varying slightly between dyads. For coherence computations, signals were smoothed at the minimum value of 12 scales.

The frequency band of interest was computed empirically rather than tying it directly to task duration as opted by previous studies ^2,7–9,11^. Figure 3 displays a coherence power spectrum plotting each of the 395 included ROIs against the 75 extracted frequency bands, averaging over all 30 dyads and all time-points. By further averaging this power spectrum over the ROIs an overall mean coherence value was obtained for each extracted scale; plotted as a red graph in figure 3. Frequencies above 220 mHz and below 9.6 mHz were removed as sampling and WTC computations introduce artifacts at the extrema. A clear positive curve indicating task related synchronicity is visible, peaking at 137 mHz. A frequency band of interest was selected based on the absolute peak differentials of the curve’s slopes, corresponding to a frequency band from 80 to 150 mHz (periods of 6.5 to 12.3 s). This band closely resembles the range selected in previous studies ^2,7–9,11^. The larger lower bound period compared to similar fNIRS-based studies was likely the result of the forced 5 second “planning”-phase initiating each trial in the current paradigm.

**Figure 3.**
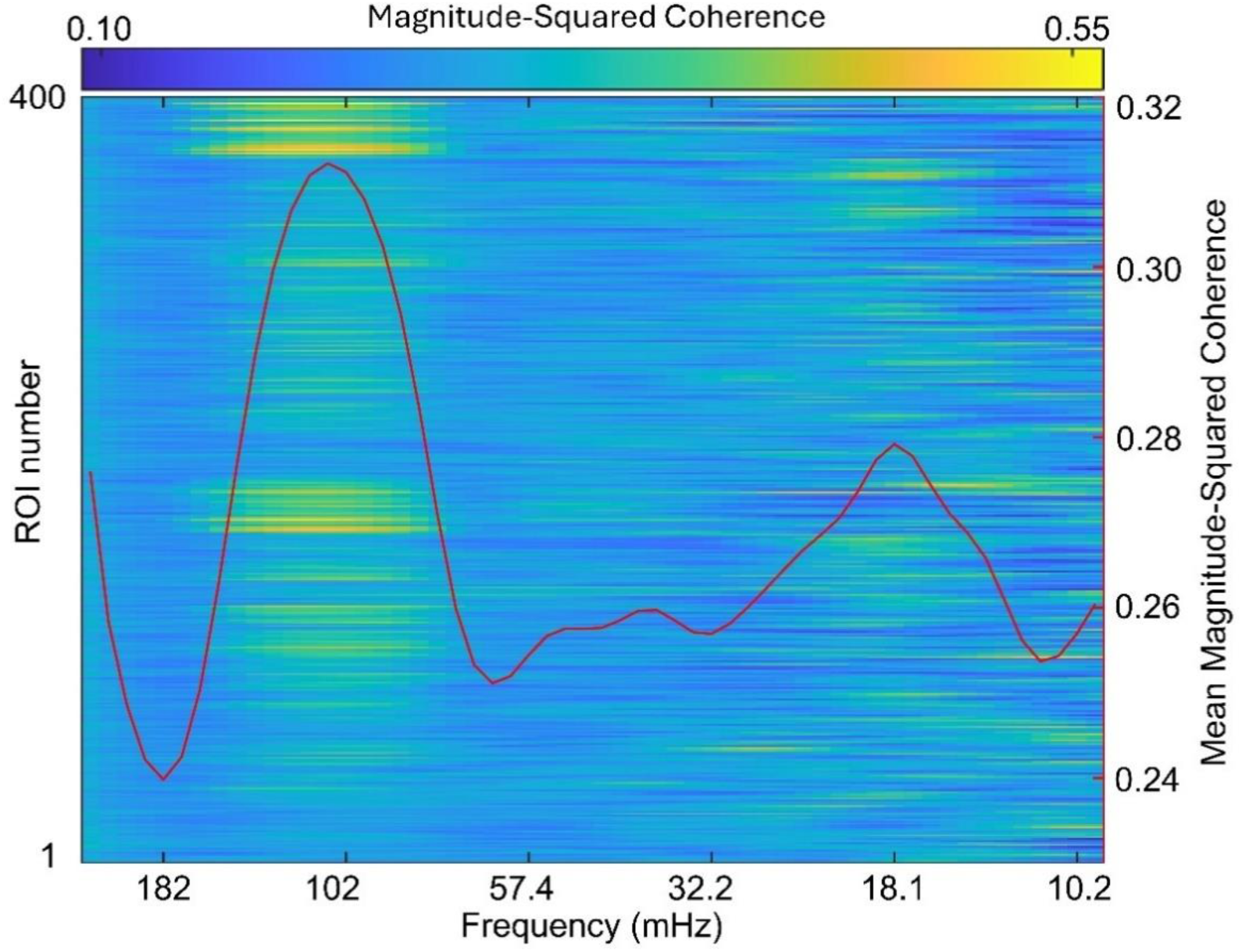
Frequency spectrum of magnitude-squared coherence of signals extracted from matching ROIs of each dyad. Plotted are frequencies between 230 and 9.6 mHz. Values are averaged over all dyads and tasks. Each row corresponds to a single ROI. A red line-plot visualizes the average of all ROIs. A positive peak is seen at 137 mHz; the frequency band of interest was empirically defined by the peak differentials of this curve, ranging from 153 to 81 mHz (periods ranging from 6.5 to 12.3 s), matching the temporal resolution of the tasks performed.

Coherence values within the selected frequency band were extracted from each task block based on the onset of the first trial and offset of the last trial. Guided by the peak of the canonical hemodynamic response function as implemented in SPM12, data obtained during the first 5 seconds of each block were excluded to account for hemodynamic delays; signal after block offset was not extracted to avoid sampling unwanted end-of-block effects. Values were then averaged over time, frequency, and session yielding one coherence term per dyad per block per ROI. Measures of coherence during the “solo” and “watch” tasks cannot be uniquely defined as the other half of the dyad was always presented with the opposing condition, values were therefore averaged to one overarching solo condition.

Our aim was to investigate task dependent changes in functional inter-brain coherence specific to interacting dyads. To correct for general task effects, a baseline coherence value was computed and subtracted from the coherence terms of each dyad. This baseline was defined as the mean of 29 permutations of the WTC procedure. For each iteration the subject ID of all subjects measured on a specific MRI scanner was manipulated, generating artificial “fake” dyads. The WTC procedure described above was then repeated for these artificial dyads. The resulting subject specific baseline describes the averaged coherence term per condition of a specific subject with each other subject assigned to the second MRI scanner. To avoid introducing a scanner specific bias, artificial dyads were never formed with two subjects scanned using the same MRI scanner.

### Behavioral data

Due to incomplete data, 3 dyads were excluded from the behavioral analysis. Voice recordings, obtained during the communication task, were manually transcribed to English text by native or professionally proficient speakers, and categorized based on their content. The category “Silence” describes trials during which neither participant spoke. “Strategic” describes trials during which task-relevant communication took place, for example: “*Press on 3*” or “*I will press faster next time*”. The category “Chit-chat” describes trials during which other, non-task relevant, forms of communication took place, for example, “*Lunch later?*” or “*Hi, how are you?*”. These categories are dummy coded in the variable “speech”. Both the number of occurrences per session and the accuracy, defined as the proportion of trials with a reaction time difference below the margin of error, were calculated per category for each dyad.

Hierarchical clustering was performed in SPSS Version 28.0 ^35^ using Ward’s method, grouping participants based on both accuracy and communication style. Differences were analyzed using squared Euclidean distance. To optimize cluster selection, the distance coefficients and associated cost (cluster count) were both standardized (range 0-1) and compared. For validation purposes, the clustering was repeated using Chi-squared measures, as the *number of occurrences* is not an interval scale variable. Both approaches yielded the same 3 large clusters with identical groupings for all pairs. To interpret the clusters, the number of trials per speech condition and accuracy of each cluster were analyzed. Mixed models with repeated measures were run in SPSS, as these account for missing data resulting from the speech labeling. Models included the factors cluster, speech, session, and the interaction term cluster*speech. Post-hoc pairwise comparisons were Bonferroni corrected.

### Personality profiles

Before the MRI investigation, participants completed several questionnaires intended to quantify their personality profiles, focusing on cooperation and/or leadership. These personality profiles improve interpretation of neuronal and behavioral findings. Specifically, participants completed the NEO Five-Factory Inventory (NEO-FFI) ^36^; the Rank Style with Pears Questionnaire (RSPQ) ^37^; the Dominance, Prestige and Leadership account (DOPL) ^38^; and the Personal Sense of Power scale (PSP) ^39^. For analyses, each factor or sub-scale was analyzed independently (NEO-FFI: Agreeableness, Conscientiousness, Neuroticism, Extraversion, and Openness to Experience; RSPQ: Dominant Leadership, Coalition-Building, and Ruthless Self-Advancement; DOPL: Dominance, Prestige, and Leadership). As any deviations from normality were relatively minor (all absolute skewness and kurtosis values below 1 and 2, respectively), linear models were run using SPSS, comparing the ratings of each subscale between clusters assigned to each participant individually. Pearson’s correlations were used to clarify the relationship between several questionnaire sub-scales. Post-hoc pairwise comparisons were Bonferroni corrected.

### Statistical models

After baseline correction, coherence values were Fisher z-transformed. Outliers were removed initially following a 1.5 inter quartile range criterion. Normality was then verified using a Shapiro-Wilk test, additionally yielding skewness and kurtosis statistics for each condition. Extrema were removed until both statistics were below critical values.

A repeated measures ANOVA was run per ROI using MATLAB 2023a, testing for differences in mean coherence values between the four conditions. Sphericity was verified using Mauchly’s tests. Greenhouse-Geisser corrections were applied for tests where sphericity was violated. Multiple comparisons over the ROIs were controlled for at a false discovery rate (FDR) of 0.05. FDR corrections were applied following the Benjamini-Hochberg procedure ^40^ using the *mafdr()* function as implemented in MATLAB. Post-hoc two-tailed paired t-test were performed for ROIs showing significant differences in mean coherence between conditions. Post-hoc tests were limited to comparisons containing the cooperation condition as these were the contrasts of interests. T-values are reported of tests significant after FDR correction. Following the same procedure, an additional exploratory analysis was performed in which a potential interaction between task condition and cluster membership (as defined by communication style and accuracy) was investigated.

## Results

### Behavioral

Though no manipulation took place, twenty-eight participants reported that they suspected that the interaction with the other participant was, at least partially, manipulated. Interestingly, a two-proportion test showed that the proportion of participants expressing at least some level of doubt did not differ between dyads that signed up for the study together compared to randomly matched participants (z = 0.915, p = .360).

Hierarchical clustering grouped participants in 3 clusters containing 14, 12, and 4 dyads, respectively. Further analysis of these clusters, and interpretation of results, were conducted in a reserved manner as the interpretation of any results was limited by this sample size. As the sub-sample of participants who were paired with a random partner was relatively small (6 dyads), including this as a factor in the models was not feasible. Independent T-tests indicated that none of the personality inventories differed significantly between participants who signed up alone and those who signed up as pairs.

Analysis of the distribution of speech among clusters showed that each of the three clusters represented dyads preferring a specific communication style (Figure 4), indicated by the number of trials during which certain speech styles occurred. This categorization was confirmed by a significant interaction between the factors cluster and speech (F(104.66, 4) = 129.79, p < .001). Notably, this interaction was not found for trial accuracy (F(106.00, 4) = 0.885, p = .476) nor was a difference between speech levels seen (F(108.428, 2) = .378, p = .686). Instead, a significant main effect on accuracy was observed for cluster (F(147.06, 2) = 3.693, p = .027). Bonferroni corrected post-hoc tests confirmed that dyads whose communication was mostly categorized as strategic discussion (n = 4) performed worse overall than the mostly chit-chatting (n = 12) dyads (M(SE) = .424(.065) vs .630(.040), p = .024). A strong trend was also seen for a difference compared to mostly silent (n = 14) dyads (M(Standard Error) = .424(.065) vs .591(.033), p = .069)(Figure 4).

**Figure 4.**
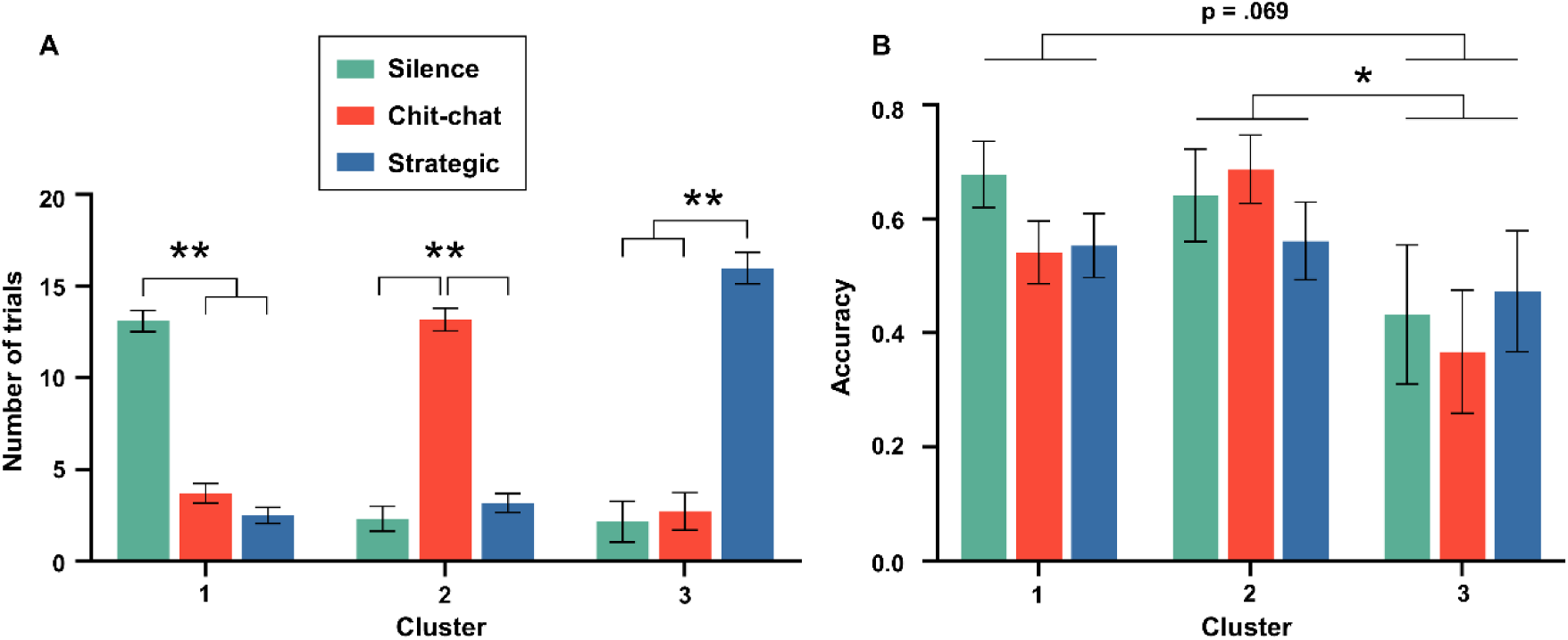
**A)** Graph visualizing the preferred communication style of each dyad-cluster. Bars depict the average number of trials per session during which a dyad engaged in a specific communication style. A strong interaction between assigned cluster and communication style is visible; each cluster presented an unambiguous preferred communication style. **B)** Graph plotting the mean task accuracy per cluster and communication style, Significant differences are seen between assigned clusters, but not communication styles. Results showed that dyads in cluster 3 performed worse during the communication task regardless of employed communication style. Error bars depict standard error. Asterisks indicate significance at p < .05 (*) and p < .001 (**)

Of all personality inventories obtained, only the neuroticism subscale of the NEO FFI and the leadership subscale of the DOPL showed significant differences between clusters (Neuroticism: F(2, 60) = 5.185, p = .008; Leadership: F(2, 60) = 3.5328, p = .036). A trend was seen for the dominant leader subscale of the RSPQ (F(2, 60) = 2.778, p = .070). Post hoc tests showed that participants who mostly chit-chatted scored significantly lower on the neuroticism than strategic participants (M(SE) = 16.08(1.483) vs 24.88(2.568), p = .013). Though chit-chatters seemed to score lower than mostly silent participants, this difference did not reach significance (M(SE) = 16.08(1.483) vs 20.64(1.373), p = 0.83). The opposite pattern was observed for the leadership subscale, here chit-chatting participants scored significantly higher than mostly silent participants (M(SE) = 25.04(1.014) vs 21.39(.937), p = .031). The trend seen for the dominant leader sub-scale of the RSPQ was mostly driven by the difference between chit-chatting and silent participants (M(SE) = 20.00(.757) vs 17.57(.701), p = .066). Neuroticism correlated negatively with both the leadership (r(60) = -.408, p = .001) and with the dominant leader (r(60) = -.399, p = .002) subscales. These two scales in turn showed a positive correlation with each other (r(60) = .724, p < .001).

### Task dependent neuronal coherence

Repeated measures ANOVAs analyzing coherence values revealed 30 ROIs in which the experimental conditions significantly (p < 0.05, FDR corrected) affected neuronal coherence (Figure 5A, supplementary Table S1). Post-hoc paired T-tests specify the origin of most of these findings (Figure 5B-D, supplementary Table S2). Where applicable, T and p ranges are provided for several neighboring ROIs covering similar regions.

**Figure 5.**
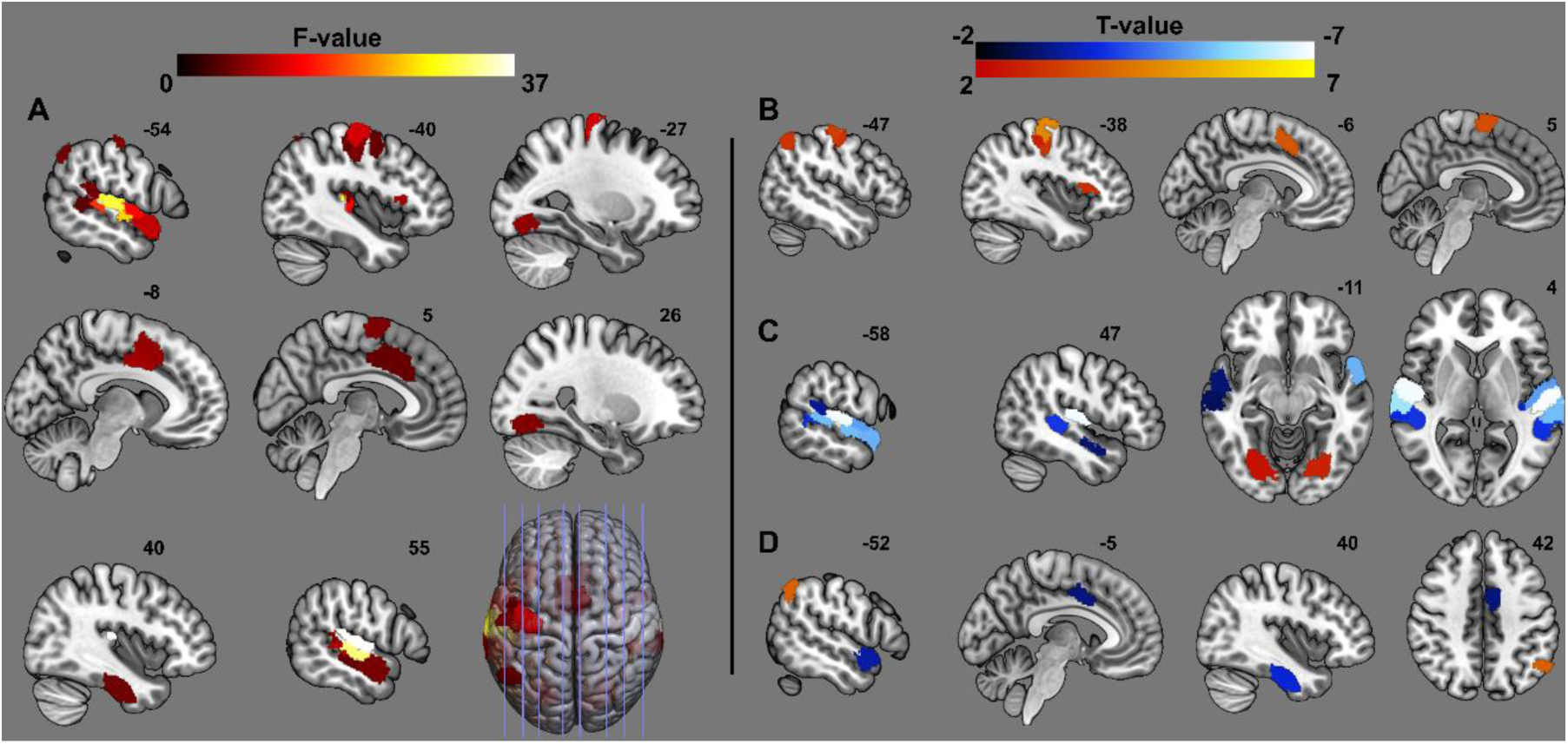
Visualization of ROIs showing task dependent differences in magnitude-squared coherence within dyads, significant at p < 0.05 FDR corrected for multiple ROIs (F-test) or multiple contrasts (t-tests). Shown are results of **A)** the repeated measures ANOVAs comparing means of each region. ROIs showing significant differences in the F-test were further investigated using paired t-tests for the contrasts **B)** cooperation - solo, **C)** cooperation - communication, and **D)** cooperation - competition.

Contrasting cooperation to solo blocks revealed significantly increased coherence in several ROIs located in left pre- and postcentral gyrus and sulcus (T range = 2.6 to 4.6, p range = .014 to < .001), left lateral occipital cortex/angular gyrus (T(25) = 3.0, p = .006), left posterior paracingulate cortex (T(27) = 3.6, p = .001), right supplementary motor area (T(26) = 3.5, p = .002), left frontal operculum cortex (T(28) = 2.9, p = .008) and left (T(29) = 3.0, p = .006) and right (T(29) = 2.7, p = .01) occipital fusiform gyrus. At the current correction levels, no ROIs showed higher coherence during solo blocks compared to cooperation tasks.

Only 2 ROIs in bilateral occipital fusiform gyrus, matching the result found of the previous contrast, showed increased coherence during cooperation compared to communication blocks (left: T = 2.7, p = .01, right: T = 2.3, p = .03). Further, this contrast was dominated by large increases in neuronal coherence for the communication task found in several ROIs in bilateral superior and middle temporal cortices, Heschl’s gyrus, planum temporale, and temporal poles (left: T range = - 3.2 to -6.9, p =< .001, right: T range = -2.7 to -6.9, p range = .01 to < .001).

Surprisingly, increases in coherence for cooperation compared to competition are limited to a single ROI in lateral occipital cortex/angular gyrus (T = 3.9, p < .001), matching the one found in the contrast against solo. This contrast did reveal several ROIs showing increased coherence for competition compared to cooperation, located in left dorsal cingulate cortex (T = -3.0, p = .006), left temporal pole (T = -3.3, p = .003), and right temporal fusiform cortex (T = -3.7, p < .001).

### Cluster-task interactions

The repeated measures ANOVA for the exploratory analysis, which included an interaction term for task condition and dyad cluster, did not reveal any ROIs in which neuronal coherence levels showed an interaction between task condition and the dyad’s assigned cluster. No further results are reported for this analysis.

## Discussion

Using a 2-scanner fMRI hyperscanning platform we were able to investigate social interaction dependent coherence of inter-brain neuronal activation while 60 participants performed an established cooperation/competition task. We showed that wavelet transform coherence is an effective way to model synchronization allowing us to fine-tune our analysis to specific temporal and frequency bands of interest. Finally, we expanded on the existing task design ^2^ by adding an additional communication condition. This also allowed us to categorize the participants based on their communication style. The use of whole-brain fMRI allowed us to localize relevant regions with greater specificity while simultaneously revealing new relevant regions only partially covered by fNIRS designs.

### Task-based neuronal coherence

Task-dependent neuronal synchronization was defined by wavelet transform coherence of empirically defined task-relevant frequency scales of the BOLD signal. By subtracting baseline coherence values, defined using coherence values of artificially paired data sets, we aimed to ensure that any remaining coherence was explained best by dyad specific interaction during the experimental blocks rather than the fact that participants performed similar tasks.

Of particular interest is the posterior paracingulate cortex (PCC) ROI which showed increased coherence during cooperation tasks. Both anterior and posterior paracingulate cortices have been strongly associated with social interaction, anticipation of behavior, and ToM ^15,41–44^, similar to the also synchronized angular gyrus ^45,46^. Stanley et al. ^47^ investigated the extent to which participants assigned human agency to an interacting entity of varying levels of humanoid or robotic actors. They suggest that the PCC and inferior parietal lobule, which in the current study also showed increased coherence during cooperation, play a role in integrating knowledge and expectations with the observed actions, especially in the case of conflicting information. In the current study, forty-two percent of participants assumed that at least some manipulation occurred during the paradigm. However, it is important to note that our current results report neuronal coherence between interacting pairs, not activation levels. In order for coherence to occur both participants should engage in similar cognitive processing; of the 30 dyads included in the fMRI analysis, only 4 had both participants express any doubt regarding the authenticity of the paradigm.

In addition to medial prefrontal cortex, superior temporal sulcus (STS) and the temporal poles are specifically relevant structures within the ToM network ^43,44^. In this context, the STS is linked to the detection of agency and goal-directed actions. While this concept is commonly studied using visually perceived action ^48–50^, our current results showed high neuronal coherence during communicated agency. While some participants questioned the authenticity of the interaction during the silent cooperation blocks, direct verbal communication with the other half of the dyad removed most if not all doubt. By communicating intentions just before each trial was performed, dyads were able to connect the performance on the task directly to the actions of another person. Within the ToM network, the temporal poles are assigned the task of retrieving cognitive scripts, described as pre-documented expectations based on a provided context ^43^. A participant’s future action on a task could be considered a script associated with the communication of one’s intent of performing this action. However, we are hesitant to describe this as the sole cause of coherence increases seen in the left temporal pole; our current paradigm was not developed to specify such high-level cognition and behavior. We must also acknowledge that some of the results seen in the temporal lobe can be explained by basic task related processes such as simultaneous auditory stimulation ^51,52^ or verbal communication ^53^, which may have survived baseline correction.

In research settings, competition is a common counterpart to cooperation as it maintains a certain level of social interaction while removing the mutual goal of the assigned task. With minor modifications, a competitive task can be employed using a similar design and stimuli. In the current study, the social aspect of the competition is effectively limited to the feedback received and how the participants process this. The optimal behavior for a motivated participant does not depend on their opponent’s behavior. During this competition task, we showed increased coherence in right dorsal cingulate cortex compared to cooperation; a region known to play a complex role in competition and reward processing. Decety et al. ^54^, however, reported the opposite pattern by showing increased activation for cooperation in left dorsal cingulate cortex. On the level of reward processing, dorsal cingulate cortex was linked specifically to reward prediction ^55^ and expectation^56^. Unifying results of competitive tasks can be challenging as their specific design can play an influential role. A popular competitive task used in both HS and single participant research is the interactive active pattern game ^54,57,58^. In the competitive variant, participants must anticipate their opponent’s moves and plan around them; this introduces strong social mentalization which we specifically aimed to minimize in our design. As mentalization was minimized for the current competitive task, finding increased neuronal coherence in left temporal pole during this task is rather surprising considering the functional relevance of this region within the ToM network. Additionally, we see that the same angular gyrus cluster found for cooperation over solo tasks, shows less coherence during competition when compared to cooperation. This suggests that the level of mentalization of the other’s behavior did differ during the two tasks, though rather than being entirely absent during competition it may have changed form. Due to the competitive nature of the task, negative feedback (“Too slow, -1 point”) can be associated with a positive outcome for the opponent which can be interpreted as a different form of mentalization. Participants were not asked how they interpreted the feedback received, preventing verification of this hypothesis.

Contrary to previous fNIRS studies we did not find significant task specific coherence differences in frontal cortices. It is possible that this is due to multiple comparison correction accounting for the large number of ROIs. Further investigation of these specific regions may thus require larger statistical power or a more limited a-priori ROI definition.

### Sensory effects

Increases in neuronal coherence in pre- and postcentral gyrus, found when comparing cooperation to solo blocks, are likely the result of key presses ^59^. While participants responded to stimuli during the solo task, their partner who was assigned to merely observe the task performed no motor actions resulting in low coherence in sensorimotor cortices for solo blocks. This task specific difference was not entirely filtered by the baseline correction as the general amplitude and consistent temporal pattern of motor activation enhance any desynchronizations found in the artificial dyads compared to the real ones. Coherence differences in bilateral occipital fusiform gyrus ROIs, found in several contrasts, best reflects high order visual areas, potentially V4 or V8, specifically sensitive for color perception specifically when consciously perceived ^60,61^. We suspect that the higher coherence levels are best explained by differing levels of attention to the color-changing stimuli during the solo task. While participants were instructed to attend to the stimuli during the watch task, the attentional demand for the spectator is, naturally, lower compared to that of a participant actively anticipating the color change and performing the task.

### Behavioral clustering

Hierarchical cluster analysis of the dyads’ performance and approach to communication revealed 3 distinct participant clusters, defined most strongly by the frequency of a specific communication style. The value of this categorization is reflected by the lack of a speech level effect on accuracy, while an effect of cluster, which is strongly driven by speech level, is observed. This implies that no single communication style directly affected accuracy significantly, rather that participant dyads that regularly employ a certain style performed worse overall. Further unraveling these effects, we see that differences in personality inventory scores between the clusters suggest that dominance and leadership may be associated with non-strategic communication while participants who focus fully on the task and limit themselves to task-relevant communication tend to score higher on the neuroticism scale. This behaviorally defined clustering did not significantly contribute to the fMRI models; this missing influence can, however, be ascribed to the limited cluster sizes.

### Limitations

As previous findings using a similar task ^7,8^ suggested that the cooperation dependent coherence was strongest for male participants, we limited our sample to this group. Naturally, this limits to what extent the findings can be generalized. Some dyads were generated by matching random participants who signed up alone. Previous research has shown positive relationship dependent effects on neuronal coherence ^9^. The sub-group of randomly matched dyads in our current sample was too small to be included as a separate factor.

As discussed earlier, cooperation, as achieved through the current task, can alternatively be described as coordination which may rely on different cognitive processes. Combined with the lacking results in frontal cortices, this limitation could suggest that the current task does not generate a social environment to the extent which the research may desire. These limitations must be taken into consideration when interpreting results in the context of social interaction. Further, any trial during which at least one participant discussed the task was labeled as “strategic”, ignoring the difference between back-and-forth communication and instructions verbalized by a single participant. Future HS studies can benefit greatly from more defined cooperation and categorized communication.

On an analysis level, comparisons of neuronal activation for each region were limited to matching anatomical locations in paired participants. As no two brains are identical, we opted for mid-sized ROIs to ensure that activation in the compared regions reflects a similar cognitive process. Consequently, this limits the spatial resolution of our analysis. Future studies may opt for a more data-driven approach through seed to voxel correlations, increasing both spatial resolution and revealing more complex inter-brain connectivity patterns. Alternatively, it may be informative to forgo ROI/seed-based analyses entirely and rather study the interactions or even coherence between cooperation associated networks ^62^. Finally, the results of the current study are discussed and compared to mainly non-HS fMRI studies. This is slightly paradoxical as the purpose of HS research is to show neural correlates of cognitive processes that are not sufficiently captured by activation levels of single participants. We hope that the field of HS-fMRI, especially in the context of synchronization and coherence, continues to expand, creating a larger scope and context for the interpretation of results.

## Conclusion and future directives

Our findings show differences in HS-related wavelet transformed coherence of neuronal activation, based on the level of cooperation within the dyads. The origin of these differences is, in part, explained by the involvement of different components of the ToM network ^41,43^, supporting previous claims ^2,8,9^. Changing the level of cooperation required during a task shifts the style of mentalization which takes place. We were pleasantly surprised by how well communication worked between participants during an fMRI measurement and highly recommend it as a tool for future HS-fMRI projects. To further unravel the cognitive mechanisms underlying social thought, mentalization, and finally behavior, it is imperative to consider all levels of interaction and communication, not just those which are convenient. Reducing social interaction in a research setting to simplified games might improve their implementation in structured scientific paradigms. At the same time, by attempting to strip such complex interactions down to what we assume are their core components, we risk losing other essential properties.

## Data availability

Scripts used to run the hyperscan synchronization server, the button pressing tasks, and analyses, as well as corrected coherence values per dyad, and data represented in the figures are available on the Open Science Framework (https://osf.io/qfx32/). Adhering to ethical restrictions, further data is available on reasonable request.

## Acknowledgments

This work was supported by the Brain Imaging Facility of the Interdisciplinary Center for Clinical Research (IZKF), part of the Faculty of Medicine, RWTH Aachen University. This project was funded by the START-Program of the Faculty of Medicine, RWTH Aachen University (Project 134/21).

We would like to thank Prof. Dr. med. Dr. rer. nat. Klaus Mathiak and Prof. Dr. med. Ferdinand Binkofski for their cooperation and high priority designation of the project, ensuring simultaneous availability of two fMRI scanners. We would also like to thank André Schüppen and Christoph Ritter for technical assistance and hardware development, Lisa Wagels for discussions and brainstorming, and finally Bernice Dauda, Julia Koch, Hayan Abo Saleh, Swati Mandekar, Samir Mammadli, Karelle Ngwen Mbang, and Paschal Ibeawuchi for support with participant recruitment and MRI measurements.

