## Supplementary Information for "Wavelet analysis of dual-fMRI-hyperscanning reveals cooperation and communication dependent effects on inter-brain neuronal coherence"

**Content:**

Tables S1 to S2

**Table S1.** Regions of interest showing significant coherence differences (F-test).

| ROI Label | Centroid Coordinates | | | F | *df* | P-value (FDR) |
|---|---|---|---|---|---|---|
| | X | Y | Z | | | |
| Left Precentral Gyrus | -35 | -18 | 64 | 11.20 | 3, 72 | < .001 |
| - | -41 | -15 | 48 | 6.22 | 3, 84 | .019 |
| - | -40 | -3 | 51 | 5.24 | 3, 66 | .040 |
| Left Postcentral Gyrus | -49 | -17 | 54 | 6.73 | 3, 78 | .041 |
| - | -39 | -25 | 53 | 5.40 | 3, 84 | .033 |
| Right Supp. Motor Cortex | 7 | -2 | 66 | 6.56 | 3, 78 | .015 |
| Left Ant. Cingulate Gyrus | -7 | 1 | 41 | 8.45 | 3, 66 | .003 |
| Left Paracingulate Gyrus | -5 | 9 | 48 | 7.71 | 3, 78 | .004 |
| Right Ant. Cingulate Gyrus | 7 | 2 | 43 | 5.13 | 3, 78 | .040 |
| Right Ant./ Paracingulate Gyrus | 7 | 19 | 35 | 5.35 | 3, 84 | .033 |
| Left Angular Gyrus/Lateral Occipital Cortex | -49 | -60 | 47 | 5.86 | 3, 66 | .025 |
| Left Heschl's Gyrus | -50 | -10 | 1 | 13.99 | 3, 69 | < .001 |
| - | -36 | -24 | 10 | 6.03 | 3, 81 | .040 |
| Left Planum Temporale | -56 | -21 | 8 | 29.62 | 3, 84 | < .001 |
| - | -59 | -37 | 16 | 6.09 | 3, 72 | .021 |
| Left Front. Operculum Cortex | -33 | 19 | 8 | 8.80 | 3, 72 | .002 |
| Left Post. Sup. Temporal Gyrus | -62 | -32 | 5 | 16.31 | 3, 87 | < .001 |
| Left anterior superior temporal gyrus | -61 | -13 | -3 | 25.05 | 3, 72 | < .001 |
| Left Mid. Temporal Gyrus | -52 | -43 | 5 | 4.91 | 3, 75 | .048 |
| Left Temporal Pole | -53 | 6 | -12 | 10.21 | 3, 63 | .003 |
| Right Heschl's gyrus | 53 | -14 | 6 | 36.61 | 3, 78 | < .001 |
| Right Planum Temporale | 60 | -24 | 11 | 36.11 | 3, 78 | < .001 |
| Right Post. Sup. Temporal Gyrus | 62 | -19 | 0 | 31.69 | 3, 87 | < .001 |
| - | 51 | -33 | 2 | 9.59 | 3, 84 | < .001 |
| - | 64 | -34 | 11 | 6.12 | 3, 87 | .019 |
| Right Ant. Sup. Temporal Gyrus | 55 | -4 | -14 | 6.17 | 3, 84 | .019 |
| Right Post. Mid. Temporal Gyrus | 63 | -23 | -7 | 5.14 | 3, 75 | .040 |
| Right Post. Temporal Fusiform Cortex | 39 | -15 | -31 | 5.81 | 3, 84 | .023 |
| Left Occipital Fusiform Gyrus | -23 | -73 | -10 | 7.65 | 3, 87 | .023 |
| Right Occipital Fusiform Gyrus | 23 | -74 | -11 | 6.72 | 3, 87 | .028 |

**Table S2.** Regions of interest showing significant contrast-specific coherence differences compared to cooperation (post-hoc paired T-test).

| **Contrast** ROI Label | Centroid Coordinates X | Y | Z | T | *df* | P-value (FDR) |
|---|---|---|---|---|---|---|
| **Cooperation > Solo** | | | | | | |
| Left Precentral Gyrus | -35 | -18 | 64 | 4.62 | 25 | < .001 |
| - | -41 | -15 | 48 | 2.96 | 29 | .018 |
| Left Postcentral Gyrus | -49 | -17 | 54 | 3.19 | 28 | .010 |
| - | -39 | -25 | 53 | 2.61 | 28 | .043 |
| Left Paracingulate Gyrus | -5 | 9 | 48 | 3.64 | 27 | .003 |
| Left Angular Gyrus/Lateral Occipital Cortex | -49 | -60 | 47 | 3.00 | 25 | .009 |
| Right Supp. Motor Cortex | 7 | -2 | 66 | 3.50 | 26 | .005 |
| Left Front. Operculum Cortex | -33 | 19 | 8 | 2.87 | 28 | .023 |
| Left Occipital Fusiform Gyrus | -23 | -73 | -10 | 2.99 | 29 | .017 |
| Right Occipital Fusiform Gyrus | 23 | -74 | -11 | 2.69 | 29 | .036 |
| **Cooperation > Competition** | | | | | | |
| Left Angular Gyrus/Lateral Occipital Cortex | -49 | -60 | 47 | 3.86 | 25 | .002 |
| **Competition > Cooperation** | | | | | | |
| Left Ant. Cingulate Gyrus | -7 | 1 | 41 | -2.99 | 25 | .019 |
| Right Post. Temporal Fusiform Cortex | 39 | -15 | -31 | -3.71 | 29 | .003 |
| Left Temporal Pole | -53 | 6 | -12 | -3.29 | 23 | .005 |
| **Cooperation > Communication** | | | | | | |
| Left Occipital Fusiform Gyrus | -23 | -73 | -10 | 2.66 | 29 | .019 |
| Right Occipital Fusiform Gyrus | 23 | -74 | -11 | 2.33 | 29 | .040 |
| **Communication > Cooperation** | | | | | | |
| Left Planum Temporale | -56 | -21 | 8 | -6.90 | 29 | < .001 |
| - | -59 | -37 | 16 | -3.18 | 28 | .011 |
| Left Temporal Pole | -53 | 6 | -12 | -5.30 | 27 | < .001 |
| Left Heschl's Gyrus | -50 | -10 | 1 | -5.10 | 27 | < .001 |
| - | -36 | -24 | 10 | -3.67 | 27 | .003 |
| Left Ant. Sup. Temporal Gyrus | -61 | -13 | -3 | -5.44 | 27 | < .001 |
| Left Post. Sup. Temporal Gyrus | -62 | -32 | 5 | -5.43 | 29 | < .001 |
| Left Mid. Temporal Gyrus | -52 | -43 | 5 | -3.88 | 25 | .002 |
| Right Planum Temporale | 60 | -24 | 11 | -6.87 | 27 | < .001 |
| Right Heschl's gyrus | 53 | -14 | 6 | -6.86 | 29 | < .001 |
| Right Ant. Sup. Temporal Gyrus | 55 | -4 | -14 | -2.90 | 29 | .021 |
| Right Post. Sup. Temporal Gyrus | 62 | -19 | 0 | -6.04 | 29 | < .001 |
| - | 51 | -33 | 2 | -3.97 | 29 | .001 |
| - | 64 | -34 | 11 | -3.74 | 29 | .002 |
| Right Post. Mid. Temporal Gyrus | 63 | -23 | -7 | -2.73 | 26 | .034 |
